# ROTS 2.0: A reproducibility-driven framework for robust statistical modeling across diverse high-throughput omics study designs

**DOI:** 10.64898/2026.06.01.729164

**Authors:** Tomi Suomi, Jalmari Kettunen, Taneli Pusa, Laura L. Elo

**Affiliations:** Turku Bioscience Centre, University of Turku and Åbo Akademi University, Turku, Finland; Institute of Biomedicine, University of Turku, Turku, Finland

**Keywords:** reproducibility, high-throughput omics, differential expression, feature ranking, biomarker discovery, software

## Abstract

Reproducibility is fundamental to reliable scientific discoveries. The reproducibility-optimized test statistic (ROTS) is a robust framework designed to identify reproducible features (e.g. genes or proteins) in high-dimensional differential expression analyses such as transcriptomics and proteomics. This is achieved by optimizing the reproducibility of feature rankings under resampling. While originally implemented for univariate settings, ROTS now accommodates multi-group comparisons, survival analysis, linear models, and linear mixed-effects models, broadening its applicability to more complex and clinically relevant experimental designs. Using diverse simulations, benchmark datasets, and real-world case studies, we demonstrate the benefits of ROTS reproducibility optimization compared to the corresponding conventional test statistics. Additionally, we illustrate the utility of the reproducibility characteristics in assessing the overall reliability of the results. To facilitate widespread adoption, ROTS is provided as an open-source software package available through R/Bioconductor. Furthermore, to broaden the user base, we now also provide a Python interface available at pypi.org/project/PyROTS/.

## Introduction

Differential expression analysis is a fundamental tool in omics research, enabling for instance the identification of potential biomarkers, disease mechanisms, and regulatory pathways by comparing molecular features (e.g. genes or proteins) across conditions. This can provide important insights into biological variation, detect disease-associated changes, and guide therapeutic developments. However, several challenges complicate the application of these techniques. These include the high dimensionality of omics data, where the number of measured features often far exceeds the sample size, requiring approaches to control false discovery rates and improve statistical power. Technical and batch effects can introduce unwanted variation, confounding the true biological signal, while sample heterogeneity can further obscure meaningful signals. Additionally, sparsity of the data and technical noise, for instance in single-cell and proteomics studies, can make it difficult to distinguish true differential expression from random fluctuations. Overcoming these challenges requires robust statistical frameworks and careful experimental design to ensure reliable and biologically meaningful results.

In its simplest form, two-group differential expression analysis can be performed using classical approaches such as Student’s t-test and Wilcoxon rank-sum test. However, they often lack power in high-dimensional settings when sample sizes are limited. Therefore, omics studies commonly use methods such as DESeq2 ^1^, edgeR ^2^, limma ^3^, and ROTS ^4^, which are widely applied to RNA-sequencing (RNA-seq) and other high-throughput omics data. These methods use different strategies in an effort to improve robustness with small numbers of samples, including moderated statistics, empirical Bayes approaches, and reproducibility optimization. For multi-group analysis of omics data, classical approaches such as ANOVA and its non-parametric alternatives can be used. Moderated approaches such as ANOVA-like models of limma ^3^ similarly allow comparisons across multiple conditions, while accounting for sample heterogeneity. In survival analysis, methods such as Cox proportional hazards models are frequently used to associate molecular features with time-to-event outcomes ^5^. For longitudinal data, linear mixed-effects models, variance-component models, and generalized estimating equations are employed to model time-dependent changes while accounting for relevant covariates and within-subject correlations. Specialized omics methods, such as MaSigPro ^6^ and RolDE ^7^, have also been developed for longitudinal designs.

Reproducibility of results is central to reliable omics research, supporting the validation of findings and their translation into meaningful biological and clinical applications. Reproducibility remains a major concern across scientific disciplines and poses significant challenges for high-throughput omics studies, where results may often fail to replicate across independent datasets or cohorts. This can stem from various factors, such as batch effects, technical biases, or small sample sizes with inadequate statistical power ^8,9^. To improve reproducibility, robust statistical approaches together with rigorous quality control are needed to identify the most promising candidates. Selecting features that are less likely to be driven by noise or data-specific artifacts helps ensure that downstream experimental validation efforts focus on biologically relevant findings, ultimately increasing the impact of omics research.

The reproducibility-optimized test statistic (ROTS) ^4^ adjusts the test statistic according to the underlying data and provides a robust and reproducible ranking of features based on their evidence for differential expression. ROTS has been successfully applied in a range of omics studies and has shown competitive performance compared to other state-of-the-art methods across data types, including bulk RNA-seq data ^10^, single-cell RNA-seq data ^11^, label-free mass spectrometry proteomics data ^12^, as well as DNA methylation ^13^ and chromatin sequencing data ^14^. Moreover, ROTS has demonstrated strong performance in independent benchmark studies covering different omics technologies ^15–20^.

Despite these successes, ROTS has yet to fully address the growing demand for more complex analyses in omics studies. To bridge this gap, we introduce here an enhanced version of the ROTS R package that implements several new capabilities to support these advanced analyses. Specifically, in addition to the modified t-statistic for two-group comparisons, we have integrated multi-group analysis using a modified F-statistic, time-to-event analysis based on Cox proportional hazards regression, and linear and linear mixed-effects modeling for more complex study designs, all utilizing the reproducibility optimization approach. We demonstrate that the ROTS framework performs consistently well when compared to the corresponding conventional statistics in identifying differentially expressed features across various types of omics data, using simulated *in silico* data, spike-in and mixture-based gold standard datasets with known differences, and publicly available real-world datasets as case studies. To facilitate widespread adoption, ROTS is provided as an open-source R package on Bioconductor. Additionally, to ensure broad availability to the scientific community, we now also provide a Python interface. This allows users to implement their workflows within Python while still using the same backend, facilitating adoption and ensuring reproducibility. The Python version is available at https://pypi.org/project/PyROTS/.

## Results

### Data-driven optimization reflects the quality of the results

To showcase the general reproducibility optimization process (**Fig. 1a**, see Methods for details), we utilized a label-free mass spectrometry proteomics benchmark dataset consisting of a tri-organism mixture with known concentration differences between two sample groups (**Fig. 1b**) ^21^. The normalized data showed expected fold changes for the measured proteins (**Fig. 1c**), that is, a 2-fold change for yeast and a 4-fold change for *E. coli* proteins. The highest reproducibility with ROTS was achieved when the top list size matched the total number of proteins that were known to differ between the sample groups (**Fig. 1d**, dashed vertical line). The smaller reproducibility peak at the beginning reflects the number of *E. coli* proteins identified, which have the larger 4-fold change. This supports the reproducibility optimization concept, where the statistical parameters are selected based on the top list size that maximizes the reproducibility over bootstrap datasets.

**Figure 1:**
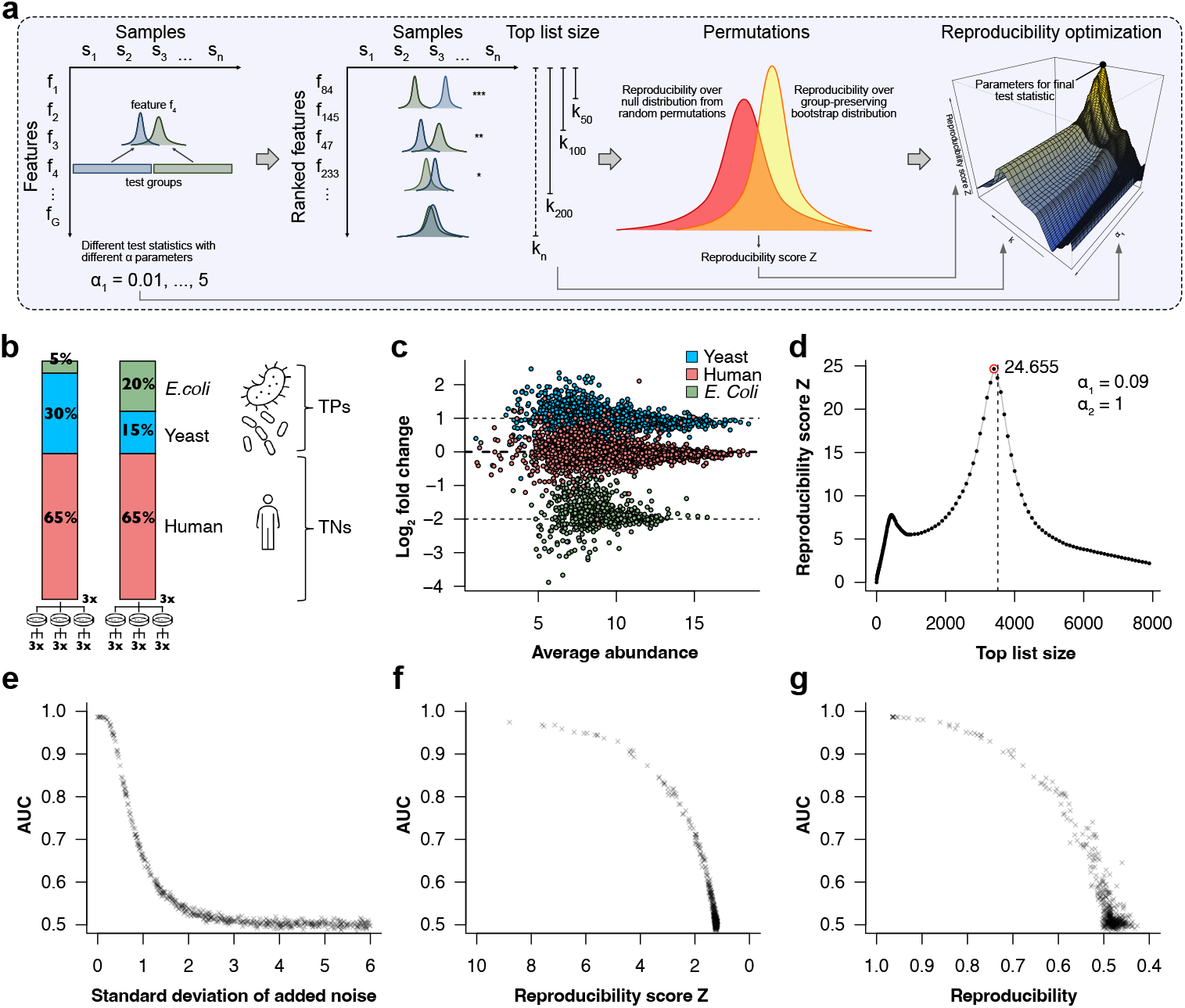
Reproducibility optimization procedure of ROTS and data-driven assessment of result quality. **(a)** Illustration of the reproducibility optimization process, where the reproducibility of top-ranked features over bootstrap datasets is maximized among a family of test statistics. **(b)** Composition of the label-free mass spectrometry proteomics benchmark dataset consisting of a tri-organism mixture with known concentration differences between two sample groups. **(c)** MA-plot showing the observed logarithmic fold changes against the average protein abundances in the mixture data, illustrating the expected 2-fold and 4-fold changes for the yeast and *E. coli* proteins, respectively, as indicated by the dashed lines. **(d)** Reproducibility score Z of the optimized parameter values plotted against different top list sizes in the mixture data. The number of proteins known to differ between the sample groups is indicated with the dashed vertical line. **(e)** Area under the ROC curve (AUC) as a function of standard deviation of added random noise, showing a decline in performance as noise increases. **(f)** Relationship between AUC and the reproducibility score Z as noise increases. **(g)** Relationship between AUC and the reproducibility as noise increases.

We further tested the performance of the method by artificially increasing random noise to the known ground truth data until the differential expression results became essentially random (**Fig. 1e**). Both the overall reproducibility score Z and the reproducibility itself decreased as random noise was added, which was reflected by a corresponding decline in the areas under the ROC curves (AUCs; **Fig. 1f** and **1g**, respectively). As expected, the AUCs dropped rapidly when the reproducibility score Z fell below two. This is well in line with the fact that the reproducibility score Z has a direct relationship to the normal distribution and, therefore, it can be used to assess the overall significance of the observed reproducibility (e.g. the critical value for p = 0.05 with a one-tailed test is 1.645). This reproducibility score represents a key advantage of the ROTS framework by providing a quantitative measure to assess the overall quality of the results, which is particularly important in real-world data, where no ground truth is typically available.

### Validation of ROTS analysis against standard models

To validate the extended functionality of ROTS, we first conducted technical comparisons between the ROTS implementations of the t-statistic, analysis of variance (ANOVA) F-statistic, Cox proportional hazards model, linear regression model, and linear mixed-effects model against their standard counterparts, which can be considered as special cases of ROTS obtained through fixing the optimization parameters (*α*_1_ = 0, *α*_2_ = 1). The test statistics were identical and the resulting permutation-based p-values from ROTS were highly correlated with those from the standard counterparts (**Supplementary Fig. 1**; Spearman correlation p < 10^-15^).

### Reproducibility optimization improves two- and multi-group statistical analysis

Two-group comparisons using ROTS have been extensively benchmarked by us and others across diverse omics data types, including bulk and single-cell transcriptomics ^10,11,16^, mass spectrometry proteomics ^4,12,15^, and epigenomics data ^13,14^. In all these contexts, ROTS has shown competitive performance among state-of-the-art methods. To further broaden this evaluation, we now confirmed the competitive performance of ROTS also in microbiome abundance comparisons, building on a recent benchmark study ^22^. The original comparison suggested that the best performance was achieved using conventional Wilcoxon test/ordinal regression or t-test/linear regression, while some widely used methods showed highly inconsistent performance ^22^. Notably, ROTS performed comparably to the best-performing methods when assessing the consistency of results between random partitions of each dataset and ranked as the top-performing method when assessing consistency between datasets from independent studies (**Supplementary Fig. 2**).

To benchmark reproducibility optimization with multiple sample groups, we utilized a data-independent acquisition (DIA) mass-spectrometry spike-in dataset from a recent benchmarking study ^23^, where six different data processing software were tested for protein identification and quantification across different DIA acquisition schemes. The dataset consisted of a standard mixture of 48 proteins (UPS1) spiked at eight different concentrations to an *E. coli* proteome background, each with three replicates. This allowed an objective assessment of statistical methods (**Fig. 2a**). We selected the data acquired using wide and narrow isolation windows, as these were suggested to perform generally best and are commonly used. For these, we focused on the data processed by Spectronaut, being a popular software suite. From the eight sample groups, we generated all possible combinations ranging from two to seven groups. This resulted in a total of 492 different multi-group test cases for benchmarking, with performance evaluated using areas under the ROC curves (AUCs).

**Figure 2:**
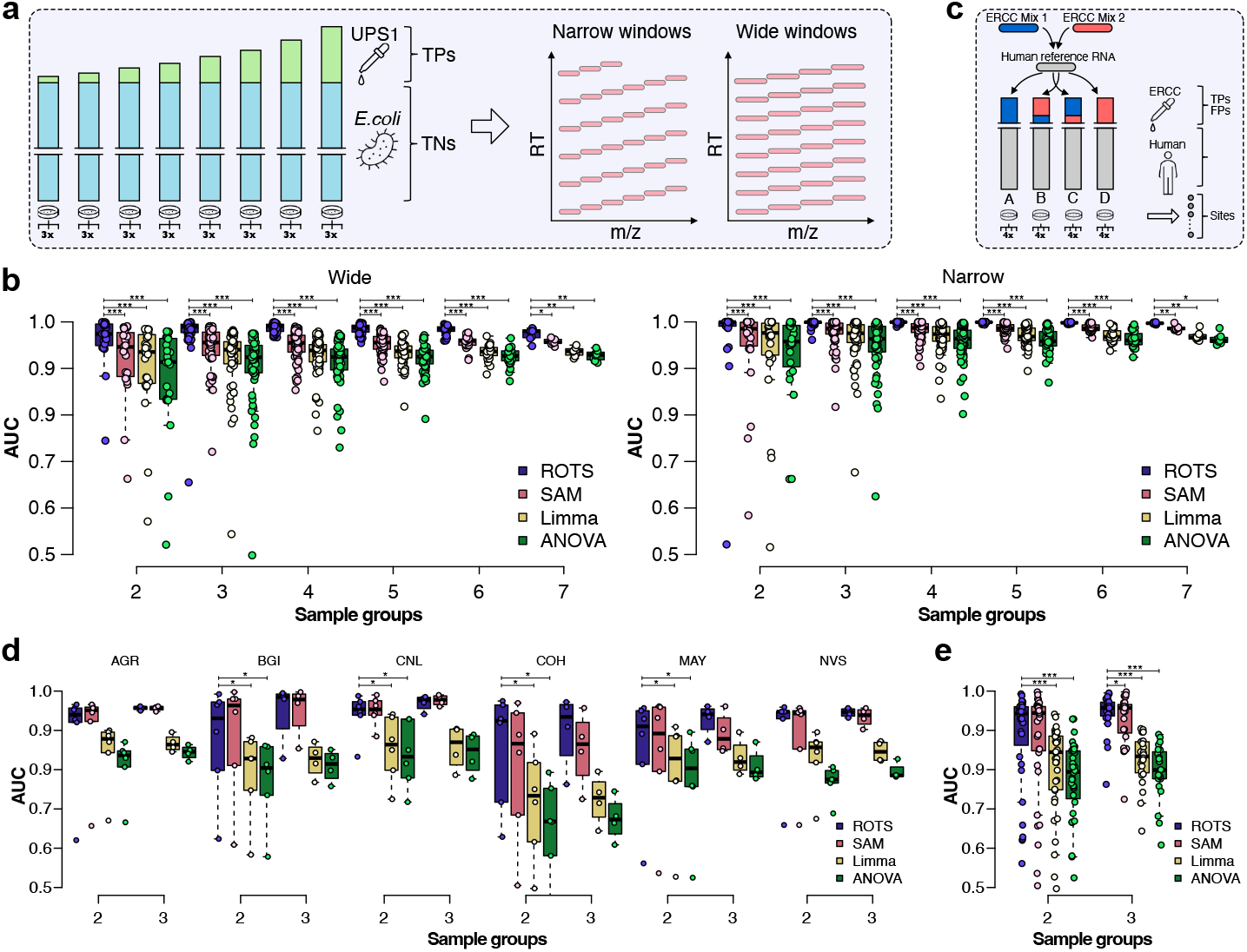
Performance of reproducibility-optimized multi-group statistical testing in spike-in mass spectrometry proteomics and RNA-seq data. **(a)** Study design of the spike-in proteomics *gold standard* dataset, where a standard mixture of 48 proteins (UPS1) was spiked at eight different concentrations to an *E. coli* proteome background, each with three replicates, and data-independent acquisition mass spectrometry was performed using both wide and narrow isolation windows. **(b)** Boxplots of areas under the receiver operating characteristic (ROC) curves (AUCs) from differential expression analyses across all possible combinations of two to seven sample groups from the wide (left panel) and narrow (right panel) isolation window proteomics data (in total 492 datasets). Wilcoxon signed rank test was used to determine statistical significance of differences in the AUC values between ROTS and other methods; *** p < 0.001; ** p < 0.01; * p < 0.05. **(c)** Study design of the SEQC RNA-seq benchmark dataset, comprising four sample types generated by adding synthetic RNA spike-in mixes to human background, each with four replicates, and sequenced across six different sites. **(d, e)** Boxplots of areas under the ROC curves (AUCs) from differential expression analyses across all possible combinations of two to three sample groups from the RNA-seq data, shown separately for each sequencing site and combined across all sequencing sites, respectively (in total 60 test cases). Wilcoxon signed rank test was used to determine statistical significance of the AUC values between ROTS and other methods; *** p < 0.001; ** p < 0.01; * p < 0.05.

The AUCs of ROTS were systematically high across the wide and narrow isolation windows as well as different numbers of sample groups (**Fig. 2b**). When compared with state-of-the-art statistical methods for multi-group analysis, including SAM, limma, and the non-optimized counterpart ANOVA, the AUCs of ROTS were significantly higher (Wilcoxon signed rank test p < 0.001; **Fig. 2b**). As expected, the performance of all methods tended to increase with the increasing number of sample groups.

We also evaluated the methods on RNA-seq data from the *Sequencing Quality Control Consortium* (SEQC) ^24^, which included four types of samples generated by adding spike-in mixes of synthetic RNA from the *External RNA Controls Consortium* (ERCC) ^25^ to a human background, each with four replicates, and sequenced across six different sites (**Fig. 2c**). Similarly as with the proteomics data, we generated all possible combinations of two to three sample groups, separately for each site, resulting in a total of 60 different multi-group test cases, and assessed performance using AUCs. This confirmed the competitive performance of ROTS, with both ROTS and SAM systematically outperforming limma and ANOVA (Wilcoxon signed rank test p < 0.05; **Fig. 2d**). Similarly, the combined assessment over all sites and group combinations supported the competitive performance of ROTS (Wilcoxon signed rank test p < 0.001; **Fig. 2e**).

### Reproducibility-optimized identification of survival-associated gene expression in TCGA breast cancer cohort

To demonstrate the use of ROTS for time-to-event analysis, we applied it to RNA-seq data from The Cancer Genome Atlas (TCGA) breast cancer cohort. Genes associated with overall survival were identified and combined into continuous cancer scores to assess their ability to capture survival-associated signals (**Fig. 3a**).

**Figure 3.**
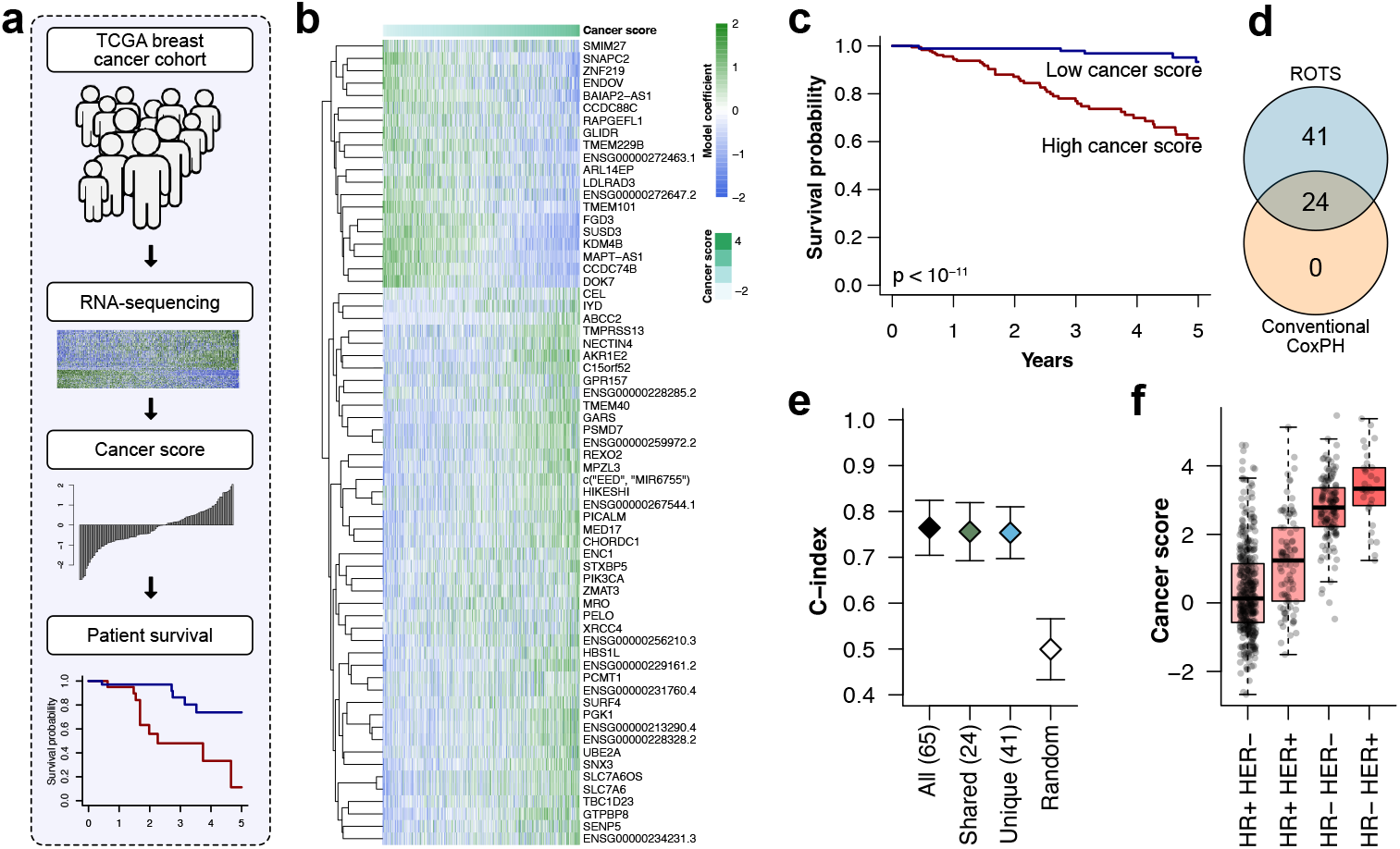
Reproducibility-optimized survival analysis in the TCGA breast cancer cohort. **(a)** Overview of the analysis workflow using RNA-seq data from the TCGA breast cancer cohort to identify genes whose expression was associated with overall survival. Genes were selected based on survival association and combined into continuous cancer scores for further evaluation. **(b)** Heatmap of genes identified by ROTS at FDR < 0.05 using Cox models with gene expression only. Patients (columns) were ordered according to the continuous cancer score. **(c)** Kaplan-Meier analysis comparing patients in the upper and lower quartile of the score distribution (log-rank test p < 10^-11^). **(d)** Venn diagram comparing genes identified using ROTS and conventional Cox regression at FDR < 0.05. **(e)** C-index values with 95% confidence intervals for cancer scores calculated using the different ROTS-derived gene sets and random gene sets of matched size. **(f)** Differences in the distribution of the ROTS-based cancer score across breast cancer subtypes based on hormone receptor (estrogen or progesterone receptor) and HER2 status (Kruskal-Wallis test p < 10^-15^).

First, we only included gene expression in the model. Using a false discovery rate (FDR) threshold of 0.05, ROTS identified 65 significant genes associated with survival. Visualization of the genes showed that patients ordered according to the cancer score showed distinct expression patterns (**Fig. 3b**). In line with this, division of the patients into high- and low-score quartiles revealed significant differences in their Kaplan-Meier curves (log-rank test p < 10^-11^; **Fig. 3c**), supporting the association of the identified genes with survival outcome.

Comparison of the significant genes identified by ROTS with those obtained using conventional Cox regression revealed 24 genes that were significant with both approaches, 41 that were unique to ROTS, and none that were unique to conventional Cox regression at FDR < 0.05 (**Fig. 3d**). To further evaluate the prognostic performance of the identified genes, we calculated C-index values for the cancer scores based on different gene sets: all 65 genes significant with ROTS, the 24 genes shared by ROTS and conventional Cox regression, the 41 genes unique to ROTS, and random gene sets of the same size as the full set of ROTS genes. Notably, C-index values of 0.7-0.8 were observed for the ROTS-derived gene sets, whereas random gene sets showed C-index values close to 0.5, as expected (**Fig. 3e**). These results supported that the genes identified by ROTS captured survival-associated information beyond that expected from random gene sets.

We also examined whether the ROTS-derived cancer score was associated with breast cancer subtypes determined by hormone receptor (estrogen or progesterone receptor) and HER2 status. As expected, significant differences were observed between the subtypes (Kruskal-Wallis test p < 10^-15^; **Fig. 3f**), with higher scores associated with clinically more aggressive subtypes. To account for potential confounding by subtype and age, we repeated the analysis using ROTS and conventional Cox models that included gene expression together with these variables. In this adjusted analysis, four genes were significant at FDR < 0.05 with ROTS, whereas conventional Cox regression did not identify any significant genes. Interestingly, the most significant gene was nectin cell adhesion molecule 4 (NECTIN4), which has previously been reported as a promising candidate prognostic marker and therapeutic target in breast cancer ^26,27^. Taken together, these results demonstrate the utility of ROTS for prioritizing robust candidate survival markers in high-dimensional omics data.

### ROTS improves linear and mixed-effects modeling in simulated omics datasets

To evaluate the benefits of reproducibility optimization for linear and mixed-effects modeling, we first compared ROTS-based analyses with their conventional counterparts using simulated datasets. The simulated datasets were designed to resemble real-world scenarios often involving multiple covariates, such as sample group, time point, individual-level effects such as sex, and technical effects such as data generation batch (**Fig. 4a**). Specifically, we simulated datasets comprising 10,000 features, each with 6-10 time points and various types of covariates. A subset of features was assigned true effects, including 150 features with differences only between two main groups, 150 features with differences between the two main groups combined with longitudinal trends, and 150 features with differences between two main groups combined with differences in a secondary group, mimicking for instance sex-specific effects. Effect sizes were simulated to range from easily detectable to very subtle that are difficult to detect, including an equal number of up- and downregulated features. Finally, additional variability was introduced through a simulated batch effect and a small systematic difference between individuals. To assess robustness, ten independent datasets were simulated using the same parameters.

**Figure 4:**
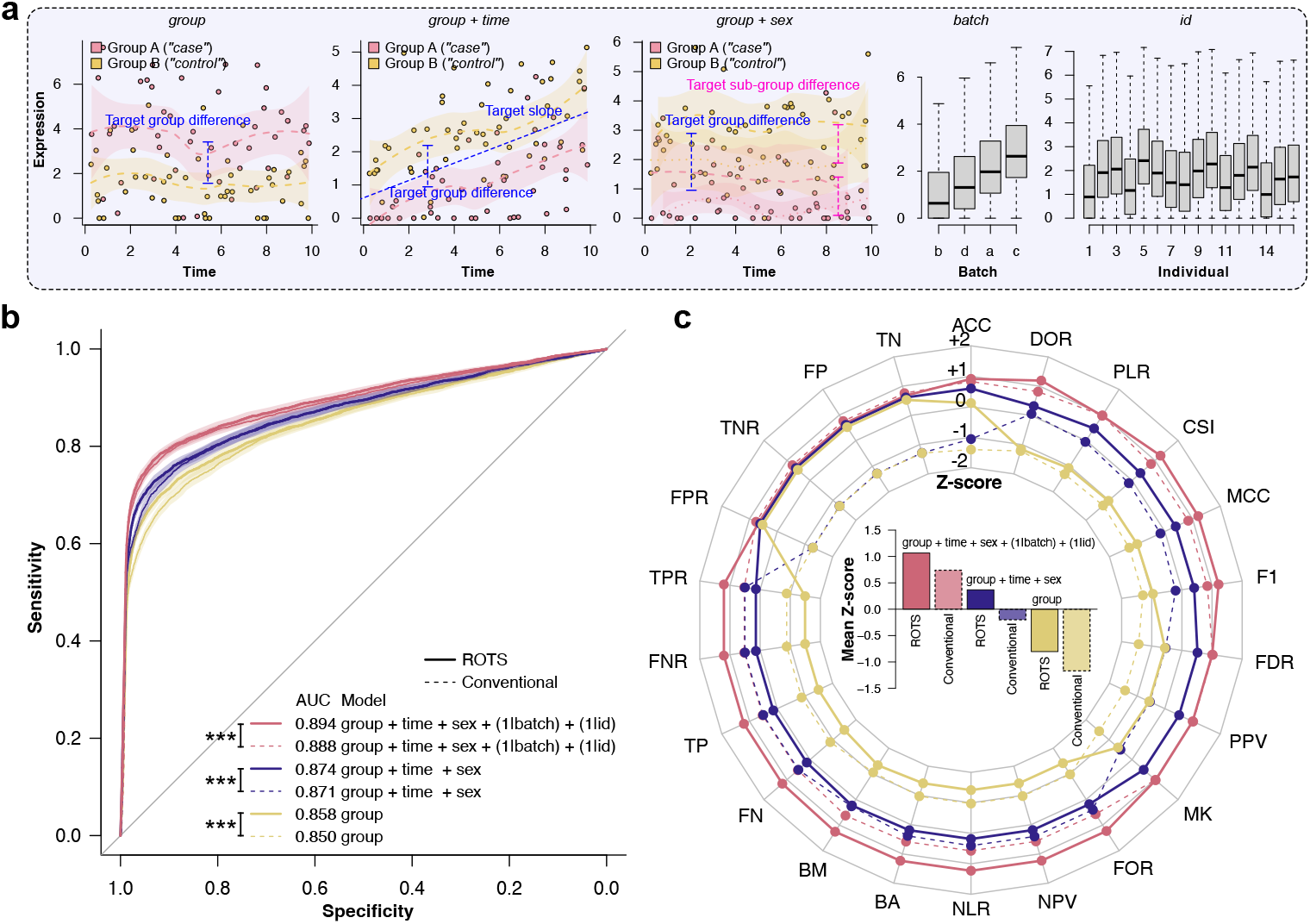
Application of reproducibility-optimized test statistic to simulated data. (**a**) Examples of simulated differences between sample groups (case and control, n=20), changes over time, subsets of data (males and females), and overall simulated batch effects and individual variation. (**b**) Receiver operating characteristic (ROC) curves for reproducibility-optimized modeling (ROTS) in comparison to conventional counterparts over ten simulations, with 95% confidence intervals, together with areas under the ROC curves (AUCs). (**c**) Performance metrics of reproducibility-optimized modeling in comparison to conventional counterparts over ten simulations, shown as Z-scores of the average value for each metric at FDR < 0.05, with positive values always indicating better performance. Metrics include the number of true positives (TP), false positives (FP), true negatives (TN), false negatives (FN), and metrics derived from them, including true positive rate (TPR), true negative rate (TNR), positive predictive value (PPV), negative predictive value (NPV), false negative rate (FNR), false positive rate (FPR), false discovery rate (FDR), false omission rate (FOR), positive likelihood ratio (PLR), negative likelihood ratio (NLR), diagnostic odds ratio (DOR), accuracy (ACC), markedness (MK), bookmaker informedness (BM), balanced accuracy (BA), F1 score, Matthews correlation coefficient (MCC), and critical success index (CSI). The barplot in the middle summarizes the values across all metrics.

To test the reproducibility-optimized modeling with the simulated datasets, three different models were considered: 1) a simple linear regression model that included only the group covariate (*group*), 2) an extended model that also incorporated additional fixed covariates (*time* and *sex*), and 3) a full mixed-effects model that additionally accounted for individual-level and batch-level random effects (*batch* and *id*). As expected, increasing the complexity of the models improved performance, as they were able to capture relevant sources of variation (**Fig. 4b-c**). In each case, the reproducibility-optimized versions provided significantly higher areas under the ROC curves than their conventional counterparts (Wilcoxon signed rank test p < 0.001; **Fig. 4b**). Similarly, performance metrics over the ten simulated datasets using a FDR cutoff of 0.05 showed that the reproducibility-optimized versions generally outperformed their conventional counterparts (**Fig. 4c**).

### ROTS linear and mixed-effects modeling identifies cross-sectional and longitudinal plasma proteomic markers of COVID-19 outcome

To demonstrate the utility of reproducibility-optimized linear modeling, we analyzed plasma proteomics data from patients with severe COVID-19 measured using Olink proximity extension assay ^28^. The dataset included 1429 plasma proteins measured in patients whose maximum 28-day acuity level was A1 (indicating death) or A2 (indicating intubation with survival). Samples were available from study days 0, 3, and 7. Patients with maximum 28-day acuity A2 were further divided by day 28 status into those who remained intubated (A2) and those who had been discharged (A5), yielding a three-level outcome for downstream analyses (**Fig. 5a**).

**Figure 5:**
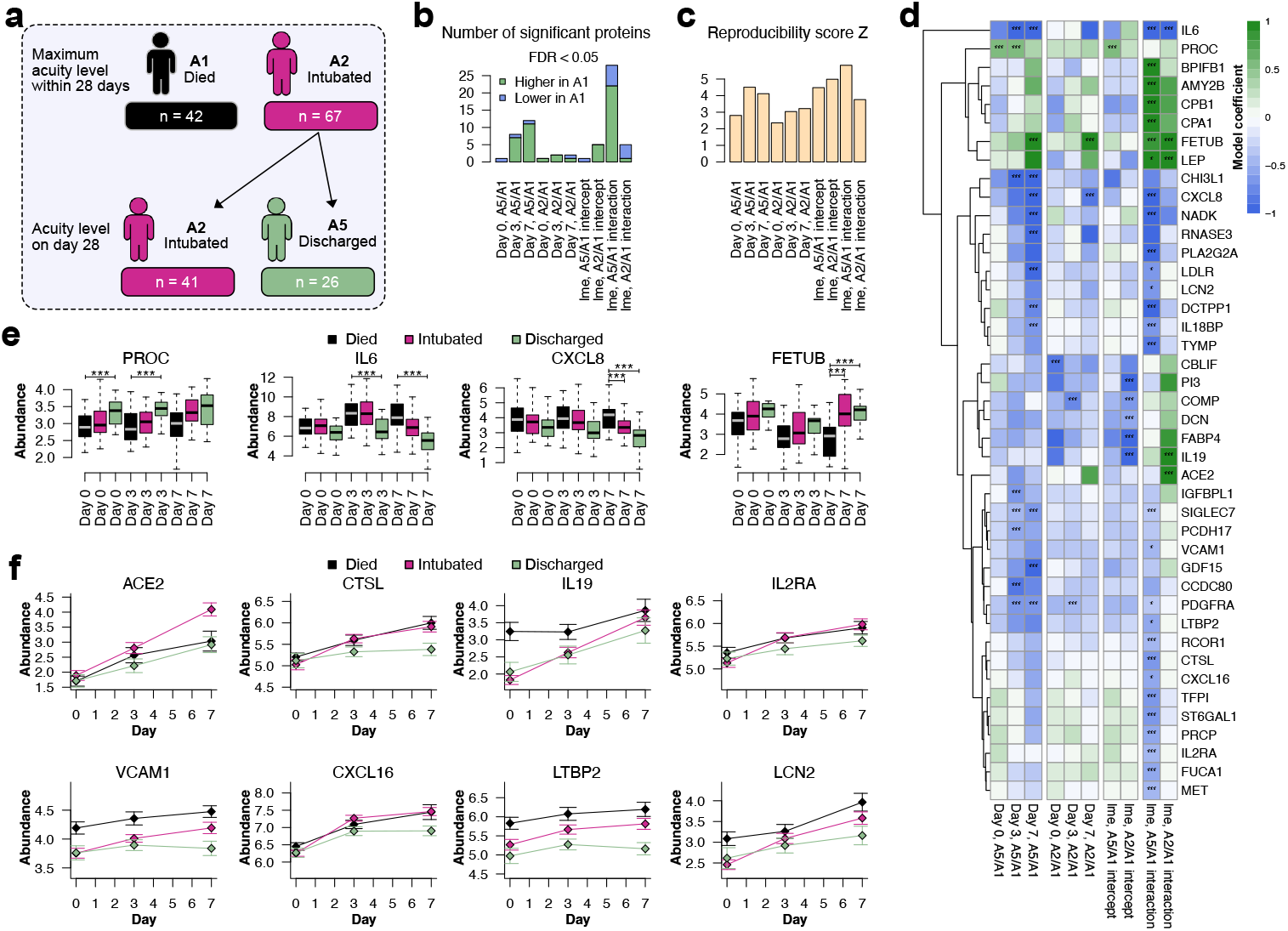
Application of reproducibility-optimized linear and linear mixed-effects modeling to real-world proteomics data from patients with severe COVID-19. **(a)** Schematic overview of the study design. The analysis included patients whose maximum 28-day acuity was A1 (died) or A2 (intubated, survived). Patients with maximum acuity A2 were further divided according to day 28 status into those who remained intubated (A2) and those who were discharged (A5), yielding a three-level outcome for the analyses. Samples were available on study day 0 (hospital admission), 3, and 7. Plasma proteomics was measured using Olink proximity extension assay. **(b)** Number of significant proteins identified using ROTS day-specific linear regression and longitudinal linear mixed-effects analyses when comparing the two survivor groups (A2 and A5) with non-survivors (A1). **(c)** Reproducibility Z-scores across the different ROTS analyses. **(d)** Heatmap of all proteins identified as significant in at least one ROTS analysis. Estimated model coefficients are shown for each comparison, with significance indicated by asterisks (* FDR < 0.05 , ** FDR < 0.01, *** FDR < 0.001). For visualization purposes, the coefficients of interaction terms were multiplied by 7 (i.e. the number of days in the data) to bring colours to a similar scale. **(e)** Representative examples of proteins identified in the day-specific ROTS analyses. Protein levels are shown separately for each outcome group and study day. **(f)** Representative examples of proteins identified through the day-by-outcome interaction in the ROTS mixed-effects modeling. Group-wise mean protein levels are shown for each study day, with error bars indicating standard errors.

We first considered day-specific linear regression models at study days 0, 3, and 7 to determine associations between protein abundance and day 28 outcome, adjusting for age and comorbidities. The number of significant proteins increased over time (**Fig. 5b**), consistent with divergence of plasma proteome profiles during severe disease. As expected, more significant proteins were identified when comparing patients discharged by day 28 with those who died than when comparing patients who remained intubated with those who died (**Fig. 5b**).

We next performed reproducibility-optimized longitudinal analysis using linear mixed-effects models, adjusting for age and comorbidities, to identify time-dependent changes in protein abundance associated with day 28 outcome. This allowed capturing dynamic signals that may not be evident in single-day comparisons. In line with the day-specific analyses, a larger number of significant day-by-outcome interaction effects was observed when comparing patients discharged by day 28 with those who died (**Fig. 5b**). In all comparisons, the ROTS reproducibility score Z was above two, supporting the robustness of the results (**Fig. 5c**).

Overall, 42 proteins were significant in at least one analysis using ROTS linear or linear mixed-effects modeling (**Fig. 5d**), including several that have previously been linked to COVID-19 severity or outcome. At day 0, the most significant association was observed for vitamin K-dependent protein C (PROC), which was also significant at day 3 and for the overall group effect in the mixed-effects model. Its abundance was lowest in patients who died (**Fig. 5e**), consistent with previous results linking low plasma levels of protein C at hospital admission to disease worsening and mortality in COVID-19 patients ^29^. At day 3, the strongest association was observed for interleukin-6 (IL6), which was also significant at day 7 and for the day-by-outcome interaction effect in the mixed-effects model. Its abundance was elevated in patients who died (**Fig. 5e**), consistent with prior studies linking elevated IL6 to mortality in critically ill patients with COVID-19 ^30^. IL6 was also among the top findings in the original study by Filbin et al. (2021) ^28^. Among proteins significant at day 7, for instance, interleukin-8 (CXCL8) and fetuin-B (FETUB) showed differences (**Fig. 5e**) consistent with previous studies, linking elevated serum CXCL8 to COVID-19 severity ^31^ and lower serum FETUB to mortality in severe COVID-19 ^32^.

Notably, a subset of proteins was identified only through the mixed-effects modeling, underscoring the added value of the longitudinal framework for detecting outcome-associated dynamic changes. Several of these proteins have previously been linked to COVID-19 severity, outcome, or pathogenesis, including ACE2 ^33^, CTSL ^34^, IL19 ^35^, IL2RA ^36^, VCAM1 ^37^, CXCL16 ^38^, LTBP2 ^39^, and LCN2 ^40^ (**Fig. 5f**).

Compared with their conventional counterparts, the ROTS day-specific and longitudinal mixed-effects modeling generally identified more significant proteins in these data, including the proteins detected by the conventional analyses (**Supplementary Fig. 3**). The exception was the day-by-outcome interaction for patients who remained intubated versus those who died, where the conventional analysis identified more significant proteins. However, the conventional approach did not detect all proteins identified by ROTS mixed-effects analysis, including IL6 and IL19. Taken together, these results show that reproducibility-optimized analysis can be applied with both linear and linear mixed-effects modeling to identify biologically relevant candidates. Although this case study was not intended to establish new COVID-19 biomarkers, the concordance of many identified proteins with prior literature supports the utility of the approach.

### GPU acceleration

The increasing scale and complexity of high-throughput omics studies have created substantial computational demands for differential expression analyses. Modern studies routinely generate datasets containing thousands to tens of thousands of features from large sample collections, making computational efficiency an important consideration. Since the ROTS framework relies on intensive resampling procedures, including permutation and bootstrapping for reproducibility optimization, computational runtime can become substantial.

To address this, we developed a GPU-accelerated implementation of ROTS that is designed to improve computational performance. Analysis of simulated data demonstrated substantial reductions in computational runtime compared with conventional CPU-based execution when increasing the number of bootstrap resamplings (**Fig. 6a**), the maximum top list size (**Fig. 6b**), the number of replicates per group (**Fig. 6c**), or the number of features in the data (**Fig. 6d**). The largest improvements in runtime were observed when increasing the number of bootstrap resamplings and the maximum top list size considered in the analysis.

**Figure 6:**
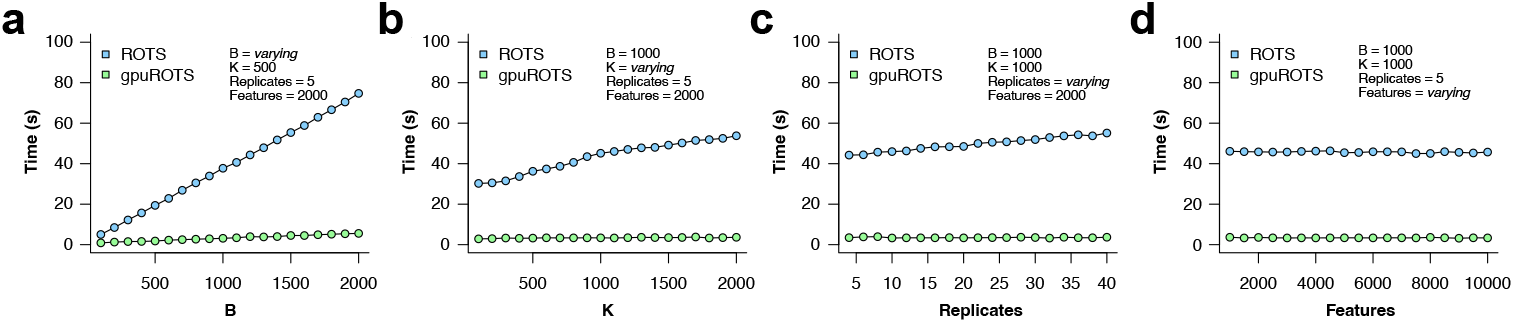
Benchmarking GPU implementation of ROTS against the conventional CPU-based implementation. Runtime comparison when increasing **(a)** the number of bootstrap resamplings (*B*), **(b)** the maximum top list size (*K*), **(c)** the number of replicates per sample group, and **(d)** the number of features (e.g. genes or proteins) in the data.

The GPU-accelerated implementation of ROTS currently focuses on classical two-group differential expression analysis, which remains one of the most commonly used settings in high-throughput omics studies. The proof-of-concept implementation is available from https://github.com/elolab/gpuROTS/.

## Discussion

Reproducibility is crucial for reliable omics discoveries, yet it remains challenging to achieve in high-dimensional omics settings where sample sizes are often limited while technical variation and biological heterogeneity can be substantial ^41–44^. In this work, we introduced an enhanced version of the ROTS R package that extends the capabilities of the reproducibility-optimized test statistic framework to support more complex study designs, which are increasingly common in omics studies, including multi-group comparisons, survival analysis, linear models, and linear mixed-effects models.

Reproducibility optimization showed competitive performance compared with conventional approaches across various simulated and benchmark datasets. Importantly, ROTS prioritizes features that consistently rank highly across resampled datasets, thereby providing a complementary approach to traditional hypothesis testing. This is motivated by the idea, closely related to stability selection, that features repeatedly prioritized across resampling are less likely to arise by random chance ^45^. ROTS also provides a measure of such robustness in terms of the reproducibility score Z which can be used to assess the overall reliability of the results. This is particularly useful in real-world omics studies where the ground truth is typically unknown.

The real-world case studies further demonstrated how the ROTS framework can be applied to more complex biological and clinical omics settings. In the TCGA breast cancer cohort, ROTS survival analysis identified candidate genes associated with overall survival. In the COVID-19 plasma proteomics data, ROTS linear and mixed-effects modeling enabled both cross-sectional and longitudinal analyses, identifying proteins associated with disease outcome while accounting for relevant covariates and repeated measurements.

A key strength of the ROTS framework is its flexibility. The method is not limited to any specific choice of effect size or variance estimation but can accommodate different options depending on user needs and be applied broadly to various statistics and estimates. By optimizing parameters within a family of test statistics based on reproducibility, ROTS can be adapted to different types of omics data and experimental designs. This flexibility does come with computational costs, as ROTS relies on repeated calculation of test statistics across bootstrapped and permuted datasets. However, this additional cost may be justified when the goal is to prioritize robust candidates for further experimental validation, which is typically substantially more time-consuming and expensive than the statistical analysis itself.

To improve computational efficiency and scalability, the ROTS implementation supports parallel computing through the BiocParallel framework in Bioconductor ^46^. This allows users to specify different parallel backends, ensuring flexibility across computing infrastructures. In addition, we introduce here a proof-of-concept GPU implementation, demonstrating that substantial runtime reductions can be achieved in computationally intensive ROTS settings, particularly when using large numbers of bootstrap resamplings or large top list sizes. More generally, GPU acceleration has emerged as a promising strategy for scaling computationally intensive omics workflows ^47–49^.

There are some limitations in this study that should be considered. First, simulations are useful for controlled benchmarking, but simulating representative high-throughput omics data is challenging because of the complexity and heterogeneity of real biological systems ^50^. Capturing realistic properties of the data is difficult, as biological variability, batch effects, and instrument-dependent technical variability should all be accurately modeled. In addition, replicating underlying biological dependencies, such as interactions, regulation through biological pathways, or temporal dynamics, adds another layer of complexity. With real data, there are also unknown biases and hidden covariates that may affect the statistical analysis. Simulations are often based on assumptions similar to those of the methods being tested, which can lead to idealized distributions that differ from real-world data ^51^. This may allow statistical models to fit the data more easily, potentially leading to overestimated performance metrics. Therefore, while simulations are useful for estimating the performance of methods and identifying possible caveats, results from real biological data should remain an important part of method evaluation ^50,52^. For this reason, we evaluated ROTS not only in simulated datasets but also using spike-in and mixture-based gold standard benchmark datasets with known differences, as well as real-world omics datasets.

Second, the current implementation of regression-based ROTS uses the same model structure for all features. This provides a consistent modeling framework across the dataset but may not capture feature-specific relationships in all settings. In some applications, researchers may be interested in considering different models for each feature, for example by adding a varying number of covariates to the model or by including quadratic terms for non-linear relationships, depending on which model fits the data best. Extending ROTS toward more flexible feature-specific model structures represents a potential direction for future development.

It is also important to recognize that general-purpose statistical frameworks like ROTS complement rather than replace domain-specific omics methods. For instance, in mass spectrometry proteomics, some existing tools can leverage peptide-level data to derive protein-level significance estimates, including MSstats ^53^, MSqRob2 ^54^, and PECA ^55^. Similarly, there are tools such as EBSEA ^56^ that use exon-level counts to detect differentially expressed genes. Such domain-specific approaches may in some cases outperform general methods applied directly to summarized counts or abundances. However, ROTS can also be incorporated into data type-specific workflows when reproducibility optimization can be applied to the corresponding statistical approach, as illustrated by its integration with PECA for peptide-level proteomics analysis ^57^.

In summary, the ROTS R package provides a reproducibility-driven approach for prioritizing robust molecular features in high-dimensional omics studies. By supporting multi-group, survival, linear, and linear mixed-effects models, it now accommodates a wide range of study designs. The open-source R/Bioconductor package, together with the newly introduced Python interface and GPU-accelerated proof-of-concept implementation, aims to make reproducibility-optimized statistical analysis accessible to a broad scientific community. We are committed to long-term maintenance of ROTS and have designed the package with user-friendliness and accessibility in mind. Its straightforward interface allows users to apply reproducibility-optimized statistical methods with minimal coding effort, while comprehensive documentation, including manuals, vignettes, and example workflows, supports both new and experienced users.

## Methods

### Reproducibility-optimized test statistics (ROTS)

ROTS optimizes the reproducibility of top-ranked features in group-preserving bootstrap datasets (i.e. resampling of the samples with replacement) among a family of modified test statistics *d*_*α*_:

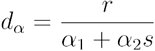

where *r* is an effect size estimate, *s* denotes a variability measure, and *α* = (*α*_1_, *α*_2_) are non-negative parameters to be optimized. Depending on the test design, the effect estimate *r* and the variability measure *s* can represent, for instance, *t*-statistic (two-group comparisons), *F*-statistic (multi-group comparisons), or regression model coefficients, as detailed in section Design-specific families of modified test statistics.

Reproducibility is defined as the average overlap of *k* top-ranked features over *B* pairs of bootstrapped datasets:

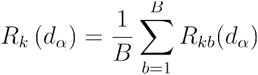

where for each pair *b* of bootstrapped datasets (*D*_*b*1_,*D*_*b*2_) the overlap is determined as

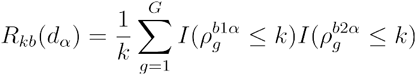

Here 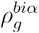 denotes the rank of feature *g* in data *D*_*bi*_ with statistic *d*_*α*_, *I* is the indicator function, and *G* is the total number of features (e.g. genes or proteins) in the data.

The optimal statistic is then determined by maximizing the reproducibility Z-score of top-ranked features at varying top list sizes *k* ∈ {1,2,…,*G*} over the parameters *α* = (*α*_1_,*α*_2_):

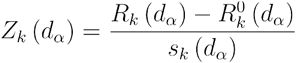

where *R*_*K*_(*d*_*α*_) is the observed reproducibility at top list size *k*, 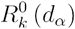 is the corresponding reproducibility in randomized datasets (permuted over samples), and *S*_*k*_(*d*_*α*_) is the standard deviation of the bootstrap distribution. The current implementation of ROTS optimizes the statistic over a dense lattice *α* = (*α*_1_, *α*_2_), where *α*_1_ ∈ {0, 0.01, …,5} and *α*_2_ ∈ {0, 1} by default.

The final ROTS output is calculated from the original data using the optimized parameters *α*_1_ and *α*_2_ corresponding to the highest reproducibility Z-score. The statistical significance and false discovery rate (FDR) are estimated by randomly permuting the sample labels. The FDR can also be determined based on the p-values using, for instance, the Benjamini-Hochberg method ^58^.

### Design-specific families of modified test statistics in ROTS

In two-group comparisons, ROTS optimizes the reproducibility of top-ranked features among a family of modified *t*-statistics, where the effect estimate *r* is the absolute difference between the group averages:

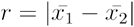

and *s* is the standard error. The current implementation of ROTS uses the pooled standard error, while the general approach is not limited to this choice:

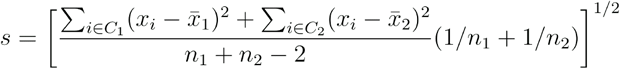

Here *i* indicates the samples in classes *C*_1_ and *C*_2_, and *n*_1_ and *n*_2_ are the number of samples in them, respectively.

In multi-group comparisons, ROTS optimizes the reproducibility of top-ranked features among a family of modified *F*-statistics, where *r* corresponds to the between-group and *s* the within-group variation:

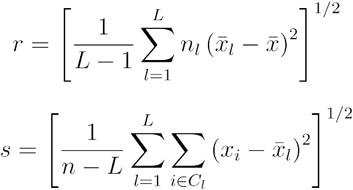

where *l* indicates the different classes {*C*_1_, …, *C*_*L*_}, *i* the samples in these classes, *n*_*l*_ the number samples in class *C*_*l*_ and *n* is the total number of samples. Here we take square roots of the between-group and within-group variance estimates, similarly as in ^59^, so that in the unmodified case the statistic is equivalent to the square root of the classical *F*-statistic.

In Cox proportional hazards regression, as well as in linear and linear mixed-effects regression analyses, the effect *r* is the estimated regression coefficient |*β*| for a variable of interest , and *s* the corresponding standard error.

### Technical validation against standard models

To validate the ROTS implementations of the t-statistic, F-statistic, Cox proportional hazards model, linear regression model, and linear mixed-effects model, we compared them against their standard counterparts. The standard versions can be obtained as special cases of ROTS by fixing the optimization parameters (*α*_1_ = 0, *α*_2_ = 1). The standard counterparts were obtained using the corresponding conventional R implementations: `t.test` for the t-statistic, `aov` for the F-statistic, `survival::coxph` for the Cox proportional hazards model, `lm` for the linear regression model, and `lmerTest::lmer` for the linear mixed-effects model. For each analysis type, we assessed the agreement of both the test statistics and the p-values between ROTS and the corresponding conventional implementation. The ROTS p-values were obtained using permutation, whereas the p-values of the conventional methods were based on their theoretical null distributions. The agreement was quantified using Spearman correlation.

### Tri-organism label-free mass spectrometry proteomics benchmark data

To illustrate the reproducibility optimization procedure, we analyzed the label-free mass spectrometry proteomics benchmark dataset from a tri-organism (human, yeast and *E. coli*) hybrid proteome mixture with known concentration differences between two sample groups ^21^. The original benchmark used different instruments and different acquisition modes. We selected here the diaPASEF data generated on a Bruker timsTOF Pro instrument, where each mixture preparation was injected in three technical replicates. The raw mass spectrometry data were downloaded from ProteomeXchange under accession PXD028735 and processed using Spectronaut (v17.7, Biognosys AG) using default search settings. Reference protein sequences for human, yeast, and *E. coli* were downloaded from UniProtKB/Swiss-Prot (2025/02). The three technical replicates were averaged for each sample, resulting in three replicate samples per group for analysis.

To assess the relationship between the reproducibility estimates and the overall reliability of the results, random noise was added to the benchmark data from a normal distribution with mean 0 and increasing standard deviation values and ROTS analysis was repeated. Performance was evaluated using the area under the receiver operating characteristic (ROC) curve (AUC), with the known differentially abundant proteins as true positives and the remaining background proteins as true negatives.

### Two-group differential abundance analysis in microbiome data

To assess the performance of ROTS in microbiome abundance comparisons, we added ROTS to the set of differential abundance analysis methods in the recent comparison study ^22^. Specifically, we applied ROTS to the split-data analyses, where each dataset was randomly divided into exploratory and validation subsets, and to the separate-study analyses, where findings from one study were evaluated in independent validation studies. All datasets, significance thresholds, and evaluation metrics were kept unchanged from the original study. For ROTS, we normalized the data using the simple total-sum scaling (TSS) normalization followed by logarithmic transformation, which is the default normalization used, for instance, in the Bioconductor package MaAsLin2 ^60^.

### Multi-group differential expression analysis in mass spectrometry proteomics spike-in data

To assess the performance of ROTS in multi-group settings, we used the data-independent acquisition mass spectrometry spike-in dataset from a recent benchmarking study ^23^. The dataset consisted of a standard mixture of 48 UPS1 proteins spiked at eight different concentrations into an *E. coli* proteome background, with three replicates per concentration. The original benchmark evaluated different acquisition window schemes and software tools. We selected here the data acquired using wide and narrow isolation windows, which were reported to perform well and are commonly used acquisition strategies, and focused on the data processed with Spectronaut (v14.10.20122, Biognosys AG).

From the eight sample groups, all possible combinations of two to seven groups were generated, resulting in a total of 492 different multi-group test cases for benchmarking. For each test case, ROTS using the modified F-statistic was compared with SAM (from R package *samr*) ^59^, limma ^3^, and the conventional ANOVA F-statistic. Performance was evaluated using the area under the ROC curve (AUC), with the known differentially abundant UPS1 proteins as true positives and the *E. coli* background proteins as true negatives. Differences in AUC values between ROTS and reference methods were assessed using the Wilcoxon signed rank test.

### Multi-group differential expression analysis in RNA-sequencing spike-in data

To assess the performance of ROTS in multi-group RNA-seq analysis, we used the Illumina RNA-seq data from the *Sequencing Quality Control Consortium* (SEQC) ^24^. The dataset included four types of samples generated by adding synthetic RNA spike-in mixes from the External RNA Controls Consortium (ERCC) ^25^ to a human background. Each sample type included four replicates and was sequenced across six sites. The raw counts were normalized using trimmed mean of M values (TMM) normalization factors estimated with the R package *edgeR* ^2^, converted to counts per million (CPM), and log_2_-transformed.

For each sequencing site, all possible combinations of two to three sample groups were generated, resulting in a total of 60 different multi-group test cases for benchmarking. For each test case, ROTS using the modified F-statistic was compared with SAM, limma, and the conventional ANOVA F-statistic. Performance was evaluated using the area under the ROC curve (AUC), with the known differentially expressed ERCC spike-in transcripts as true positives and the background genes as true negatives. Differences in AUC values between ROTS and reference methods were assessed using the Wilcoxon signed rank test.

### Survival-associated gene expression in TCGA breast cancer cohort

To demonstrate the utility of ROTS for time-to-event analysis, we analyzed RNA-seq data from The Cancer Genome Atlas (TCGA) breast cancer cohort, using overall survival as the outcome. Raw counts and clinical information were downloaded from the recount2 resource ^61^. Counts were normalized using TMM normalization factors estimated with the R package *edgeR* ^2^, converted to counts per million (CPM), and log_2_-transformed. Genes with mean logarithmic CPM < 1 across samples were filtered out. Overall survival was capped at 60 months.

Cox proportional hazards models were considered at two levels: using gene expression only, and using gene expression together with age and subtype information in the model. Subtypes were determined based on hormone receptor status, defined by estrogen receptor or progesterone receptor positivity, and HER2 status. Genes were considered significant at FDR < 0.05.

To evaluate the prognostic information captured by the significant genes, continuous cancer scores were calculated for each sample similarly as in ^62^. Briefly, gene expression values were scaled across the patients as standard deviations from the median. Within each patient profile, the score was defined as the two-sided t-statistic comparing genes associated with better and worse survival. Kaplan-Meier analyses were performed by dividing patients into upper and lower, quartile groups based on the cancer score distribution. Survival differences were assessed using the log-rank test.

Harrell’s concordance index (C-index) was calculated to compare the prognostic performance of cancer scores based on different gene sets, including all genes significant with ROTS, genes significant with both ROTS and conventional Cox regression, genes unique to ROTS, and random gene sets of the same size at the full set of ROTS genes. The C-index quantifies the proportion of comparable patient pairs for which the predicted score ordering agrees with the observed survival ordering.

### Analysis of simulated longitudinal data

Synthetic longitudinal omics datasets were generated to evaluate the performance of ROTS linear and linear mixed-effects models under controlled settings. For each dataset, we first simulated subject-specific sampling times for a desired number of individuals (n = 20), with each individual measured at six to ten time points, sampled randomly from a uniform distribution between 1 and 10, representing for instance years or months. Additional subject-level variables, including group and sex, were distributed evenly across the individuals. In addition, an artificial batch was defined by randomly assigning each sample to one of four batches.

A high-dimensional feature-by-sample matrix was then generated for 10,000 features, with baseline expression values sampled from a normal distribution with a mean of 2.5 and standard deviation of 0.5 Feature-wise noise was added with a mean of 0 and standard deviation scaled according to the underlying expression level, ranging from 0.01 to 0.5.

Biological signals were introduced into subsets of features to represent different modeling scenarios. The first 150 features contained group differences only, with effect sizes ranging from -1 to 1, representing effects from easily detectable to subtle near-zero differences including equal numbers of up- and down-regulated features. The next 150 features contained combined group and time effects, with time effect coefficients ranging from -0.75 to 0.75 in addition to group differences. These features were divided into ten different subsets with different crossover points, where the groups have equal expression level, distributed evenly from the beginning to the end of the time series. The next 150 features contained both group and sex effects, with sex effect sizes ranging from -2 to 2 in addition to the group differences. A further 150 features contained only time trends, with time effect coefficients ranging from -0.75 to 0.75, and another 150 features contained only sex effects, with effect sizes ranging from -2 to 2.

Individual-level random effects were simulated by adding individual-specific constants sampled from a normal distribution with a mean of 0 and standard deviation of 0.05. Batch effects were simulated by adding batch-specific constants ranging from -1 to 1. Negative values generated during the simulation were replaced with zero. Ten independent datasets were generated using the same simulation settings.

The simulated datasets were analyzed using three model formulations: a simple linear regression model including only the main group covariate, an extended linear model including group, time, and sex as fixed covariates, and a linear mixed-effects model, including the fixed covariates together with random effects for individual and batch. For each model formulation, the ROTS-based analysis was compared with the corresponding conventional analysis. Performance was evaluated using the area under the ROC curve (AUC), with the known true positives and true negatives. Differences in AUC values between ROTS and conventional analyses were assessed using the Wilcoxon signed rank test. Additional performance metrics were calculated at FDR < 0.05, including the number of true positives (TP), false positives (FP), true negatives (TN), false negatives (FN), and metrics derived from them, including true positive rate (TPR), true negative rate (TNR), positive predictive value (PPV), negative predictive value (NPV), false negative rate (FNR), false positive rate (FPR), false discovery rate (FDR), false omission rate (FOR), positive likelihood ratio (PLR), negative likelihood ratio (NLR), diagnostic odds ratio (DOR), accuracy (ACC), markedness (MK), bookmaker informedness (BM), balanced accuracy (BA), F1 score, Matthews correlation coefficient (MCC), critical success index (CSI), and summarized across the ten simulated datasets. For visualization, the performance metrics were transformed into z-scores so that positive values indicated better performance.

### Analysis of plasma proteomics data from patients with severe COVID-19

To demonstrate the application of ROTS linear and linear mixed-effects modeling to real-world omics data, we analyzed plasma proteomics data from patients with severe COVID-19 ^28^. The proteomics data together with clinical information were downloaded from Mendeley Data (https://data.mendeley.com/datasets/nf853r8xsj/1). The proteomics data consisted of measurements using the Olink proximity extension assay. We used the readily processed relative abundance (NPX) values, which included 1429 plasma proteins available for analysis without any missing values. Data were available from study days 0, 3, and 7. The clinical covariates available included age as a five-level categorical variable (20-34, 36-49, 50-64, 65-79, and 80+ years) and six binary comorbidity indicators (pre-existing heart disease, diabetes, hypertension, pulmonary disease, kidney disease, immuno-compromised condition). Here, we restricted the analysis to COVID-19 patients whose maximum acuity level within 28 days of enrollment was A1 (indicating death) or A2 (indicating intubation with survival), as defined in the original study ^28^. Patients with maximum 28-day acuity A2 were further divided by their day 28 status into those who remained intubated (A2) and those who had been discharged (A5), yielding a three-level outcome for the analyses.

Linear and linear mixed effects models were fit separately for each protein using the NPX values. For day-specific linear models, analyses were performed separately for study days 0, 3, and 7 using the acuity level as the main variable of interest, with A1 as the reference group, and adjusted for age and comorbidities to control for potential confounding. Age was used as a five-level factor, with the youngest age category as the reference, and comorbidities as binary covariates. For the linear mixed effects models, data from all available time points was used, with fixed effects for study day, acuity level, their interaction, age, and comorbidities, together with a random intercept for patient. Study day was modeled as a numeric variable with values 0, 3, and 7 to focus on linear temporal trends in relative protein abundance. The primary effect of interest was the interaction between study day and acuity level. Proteins were considered significant at FDR < 0.05.

### Statistical analysis

All statistical analyses were performed in R version 4.5.1.

### GPU-accelerated implementation

A proof-of-concept GPU-accelerated version of ROTS was implemented using the OpenCL framework through the *OpenCL* R package, where the computationally intensive parts are implemented as OpenCL kernels. The R interface takes care of overall functionality and runtime kernel compilation, while numerical operations are offloaded to the GPU device.

Runtime benchmarking was performed using simulated datasets with values sampled from normal distribution with mean 14 and standard deviation 1, where 5 % of rows had simulated upregulation and 5 % had simulated downregulation between sample groups. Runtime was compared between the GPU implementation and the conventional CPU-based implementation while varying the number of bootstrap resamplings from 100 to 2000, the maximum top list size from 100 to 2000, the number of replicates per sample group from 4 to 40, and the number of features (e.g. genes or proteins) in the data from 500 to 10,000. Benchmarks were performed on 3.2GHz Intel Xeon CPU and AMD Radeon Pro W5700X GPU having 2560 stream processors.

### Python interface

The Python interface, PyROTS, uses rpy2 (https://rpy2.github.io/) to provide a Python interface for the ROTS R package. This allows the user to implement their workflow entirely within Python using native data structures while running the same underlying R code for the analysis.

### Software availability

The ROTS R package is available through Bioconductor: https://bioconductor.org/packages/ROTS/ The Python interface is available through PyPI: https://pypi.org/project/PyROTS/ The GPU implementation is available through GitHub: https://github.org/elolab/gpuROTS/

## Acknowledgements

The authors wish to acknowledge CSC – IT Center for Science, Finland, for computational resources.

The results here are in part based upon data generated by the TCGA Research Network: https://www.cancer.gov/tcga.

---

**Supplementary Figure 1.**
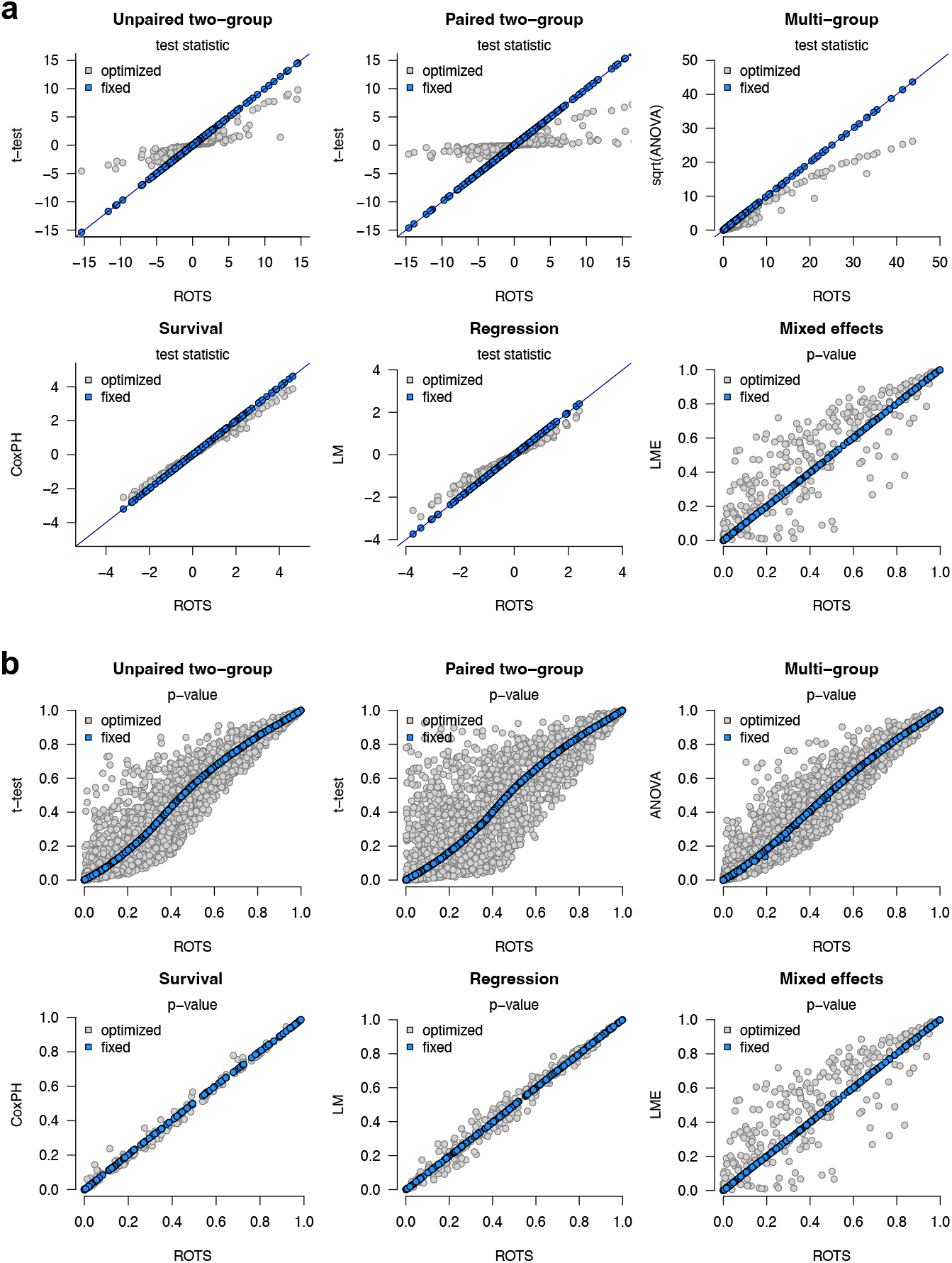
Technical comparison of ROTS implementations with their standard counterparts. Comparisons of the ROTS implementations of paired and unpaired t-statistics, analysis of variance (ANOVA) F-statistic, Cox proportional hazards model, linear regression model, and linear mixed-effects model with their standard counterparts. Two-group and multi-group tests are performed on Gotti et al. (2021) spike-in proteomics benchmark data between 10 fmol and 25 fmol injections, and by using 5, 10 and 25 fmol injections, respectively. Survival statistics are performed on Hirvonen et al. (2023) type 1 diabetes follow-up proteomics study, with time to diagnosis as the survival time. Regression model was performed on the same study, with study group and age as the explanatory variables, and where mixed effect model further included individual as random variable. **(a)** Comparison of test statistics. **(b)** Comparison of p-values.

**Supplementary Figure 2.**
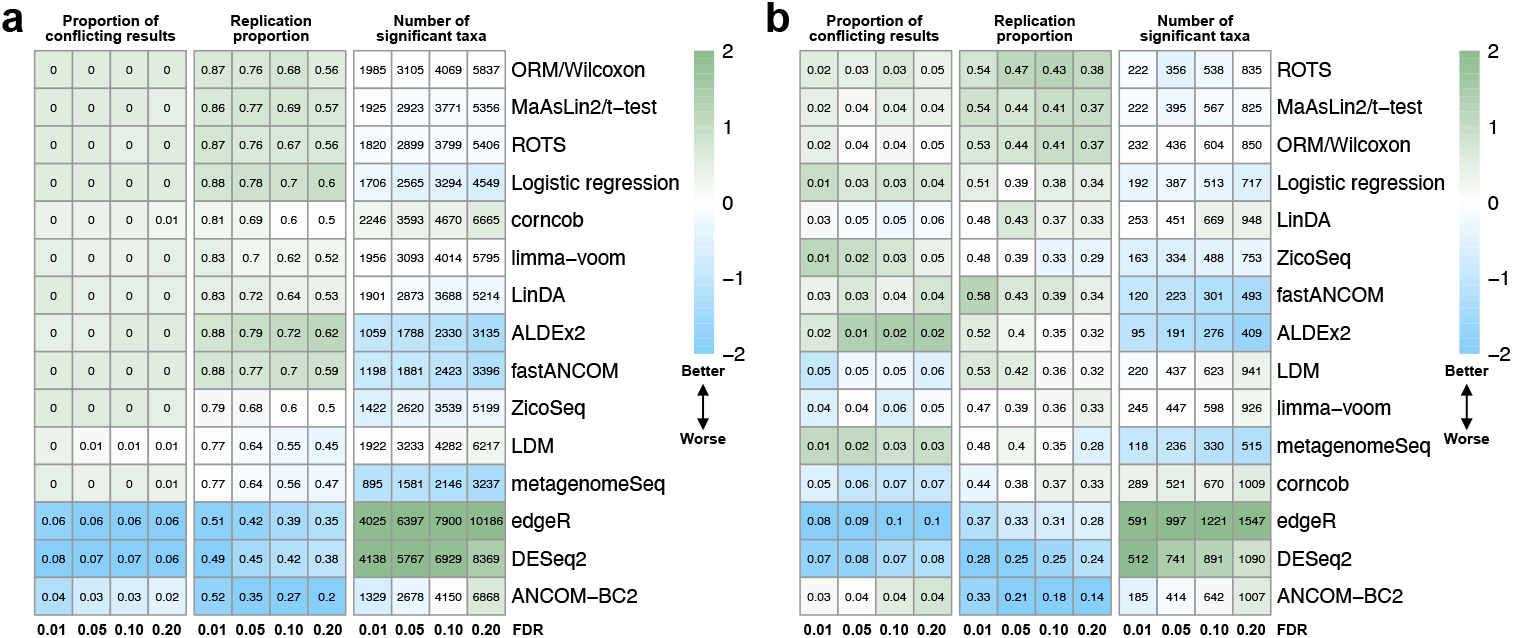
Performance of differential abundance analysis methods in microbiome data. Performance of ROTS was compared with 14 differential abundance analysis methods (rows) evaluated in a previous comparison study using taxonomic profiling data from 16S rRNA or shotgun metagenome sequencing. **(a)** Consistency of the methods in 57 randomly split microbiome datasets, with each original dataset split five times into two equally sized subsets, resulting in a total of 285 pairs of exploratory and validation datasets. **(b)** Consistency of the methods between independent studies, using 37 datasets as exploratory datasets. The proportion of conflicting results denotes the proportion of taxa that were significant at the selected FDR threshold (columns) in the exploratory dataset and also significant in the validation dataset but in the opposite direction. The replication proportion denotes the proportion of significant taxa that were significant in the same direction in both the exploratory and validation datasets. The methods were ordered based on the mean of the standardized values of the performance metrics.

**Supplementary Figure 3.**
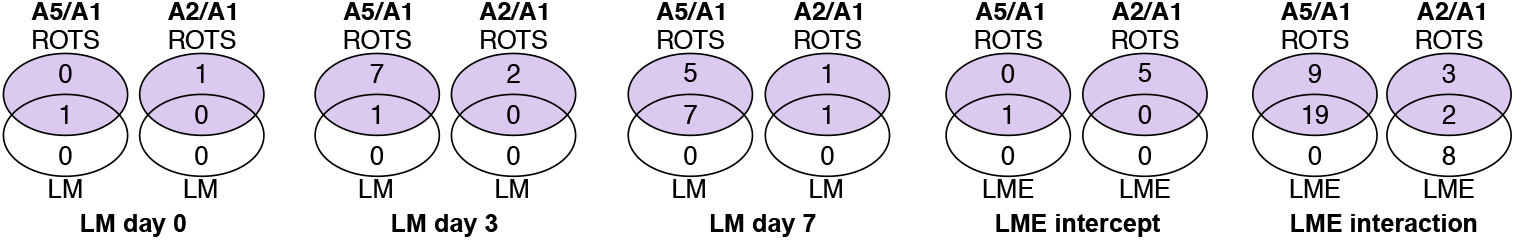
Overlaps of significant proteins identified by ROTS and conventional linear and mixed-effects models in the COVID-19 plasma proteomics data. Overlaps of significant proteins (FDR < 0.05) identified by ROTS and the corresponding conventional models in the day-specific linear regression (LM) analyses at study days 0, 3, and 7, and in the longitudinal linear mixed-effects (LME) analyses. In the mixed-effects analyses, both overall group effects (intercept) and day-by-outcome interaction effects were considered. Comparisons were performed between patients who remained intubated (A2) or were discharged (A5) and those who died (A1), adjusting for age and comorbidities.

